# A complete tool set for molecular QTL discovery and analysis

**DOI:** 10.1101/068635

**Authors:** Olivier Delaneau, Halit Ongen, Andrew Brown, Alexandre Fort, Nikolaos Panousis, Emmanouil Dermitzakis

## Abstract

Population scale studies combining genetic information with molecular phenotypes (e.g. gene expression) become a standard to dissect the effects of genetic variants onto organismal. This kind of datasets requires powerful, fast and versatile methods able to discover molecular Quantitative Trait Loci (molQTL). Here we propose such a solution, QTLtools, a modular framework that contains multiple methods to prepare the data, to discover proximal and distal molQTLs and to finally integrate them with GWAS variants and functional annotations of the genome. We demonstrate its utility by performing a complete expression QTL study in a few and easy-to-perform steps. QTLtools is open source and available at https://gtltools.github.io/gtltools/

## Main text

In order to increase the explanatory power of genome wide association studies (GWAS), many genetic studies now routinely combine genetic information with one or multiple molecular phenotypes such as gene expression [1], protein abundance [2], metabolomics [3], methylation[4] or chromatin activity [5]. This makes possible the discovery of molecular Quantitative Trait Loci (molQL); a key step towards better understanding the effects of genetic variants on the cellular machinery and eventually on organismal phenotypes. In practice, this requires analyzing datasets comprising millions of genetic variants and thousands of molecular phenotypes measured on a population scale; a design that aims to perform many orders of magnitude more association test than in a standard GWAS. To face this computational and statistical challenge, there is a clear need of computational methods that are (i) powerful to properly face the multiple testing problem, (ii)fast to easily process large amount of data in reasonable running times and (iii) versatile to adapt to new datasets as they are being generated. Here, we present such an integrated framework, called QTLtools, which allows users to go from raw sequence data to collections of molQTLs in only few easy-to-perform steps,all based on powerful methods that either match or improve those employed in large scale reference studies such as Geuvadis [1] or GTEx [6].

QTLtools is a modular framework designed to accommodate new analysis modules as they are being developed by our group or the scientific community. In its current state,QTLtools performs multipe key tasks (figure 1) such as checking the quality of the sequence data, checking that sequence and genotype data match, quantifying and stratifying individuals using molecular phenotypes,discovering proximal or distal molQTLs and integrating them with functional annotations or GWAS data.To demonstrate the utility of this new tool with real data, we used it to perform a complete expression QTL (eQTL) study for 358 European samples where genotype and expression data were generated as part of the 1000 Genomes [7] and Geuvadis [1] projects, respectively (supplementary material 1).

**Figure 1:**
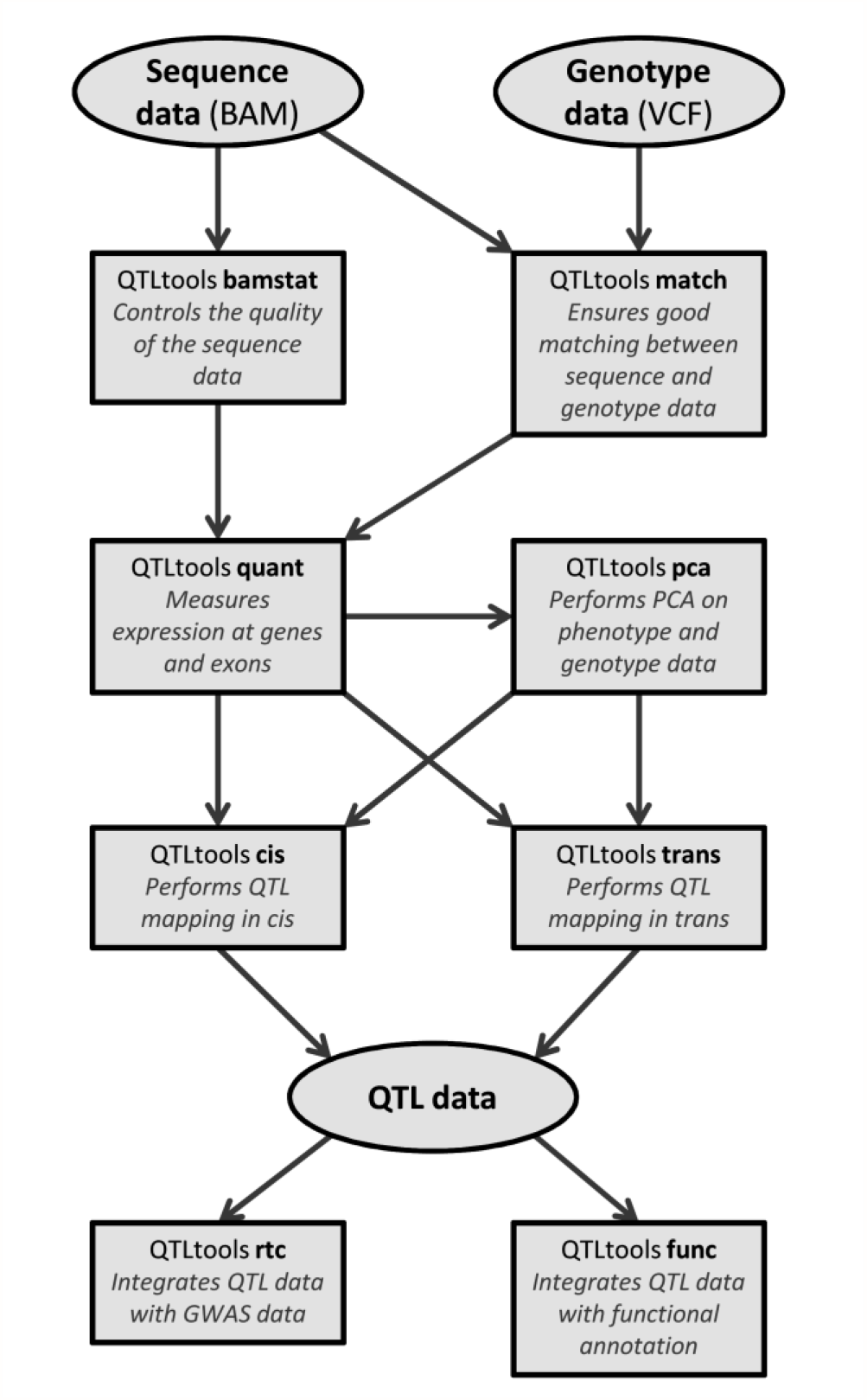
Flow chart of the main QTLtools functionalities. This represents how the various functionlities of QTLtools can be combined in order to go from the raw sequence and genotype data to collections of molecular QTLs which can then be integrated with both GWAS data and functional annotations. Data is represented with ovals and tasks with boxes in which the name of the mode is shown in bold black with a short description of what it does.

To control quality of the sequence data, QTLtools proposes two complementary approaches. First,it can measure the proportions of reads (i) mapping to the reference genome and (ii) falling within an annotation of interest (supplementary method 1), such as GENCODE for RNA-seq [8]. Second,it can also make sure that the sequence data matches the corresponding genotype data; the opposite being an evidence of sample mislabeling [9]. To achieve this, QTLtools measures concordance between genotypes and sequencing reads, separately for heterozygous and homozygous genotypes (supplementary method 2). Low values in any of the two measures indicate problems such as sample mislabeling, contamination or amplification biases (supplementary figure 1).When performed on Geuvadis, these two approaches demonstrated the high quality of the RNA-seq data and the good match with available genotype data (supplementary figures 2-3).

To quantify gene expression, QTLtools counts the number of sequencing reads overlapping a set of genomic features (e.g. exons) listed in a given annotation file (supplementary method 3).We quantified both exon and gene expression levels in all358 Geuvadis samples using this approach and get 22,147 genes quantified in more than half of the samples (supplementary figure 4). Then, we run principal component analysis(PCA) on these quantifications as implemented in QTLtool (supplymentary method 4) in order to capture any stratification in the sequence data or the genotype data. In the Geuvadis data we did not observe any unexpected cluster in the expression data, neither in the genotype data (supplementary figure 5) and used the resulting sample coordinates on the first Principal Components as latent variables to increase discovery power of any downstream association testing (supplementary method 5).

A core task of QTLtools is to discover proximal (i.e. *cis*-acting) molQTLs. To do so, it extends the QTL mapping method introduced by FastQTL [10] and offers multiple key improvements that make this step fast and easy-to-perform. First, it uses a permutation scheme that needs a relatively small number of permutations to adjust nominal p-values for multiple testing (supplementary method6, supplementary figure 6). As a consequence, the whole Geuvadis eQTL analysis can be performed in short running times (˜32 CPU hours) which has been proved to be an order of magnitude faster than a widely used tool, Matrix eQTL [11] and provides adjusted P-values without any lower bounds (supplementary figure 7). The running times are actually so small that it becomes easy to repeat the whole analysis multiple times across different sets of quantifications, covariates and QC filters in order to determine the optimal configuration maximizing the number of discoveries (supplementary figure 8-9). In addition, QTLtools also provides ways to easily extract subsets of data and therefore facilitate detailed inspection of particular eQTLs (supplementary figure 10). As multiple molecular phenotypes can belong to higher order biological entities such as exons of genes or histone modification peaks to Variable Chromatin Modules (VCMs) [2], we also implemented two methods to maximize the discoveries in such particular cases (supplementary method 7). Specifically QTLtools can either (i) aggregate multiple phenotypes in a given group into a single phenotype via PCA or (ii)directly use all individual phenotypes in an extended permutation scheme that accounts for their number and correlation structure. In our experiments, the permutation-based approach seems to outperform the PCA-based approach in term of number of discoveries in the two data sets we tested (figure 2A, supplementary figure 11). In Geuvadis, the permutation-based approach is able to discover an additional set of ˜1,056 eQTLs compared to the standard gene-level quantification,most of them being for genes containing many exons (supplementary figure 12).

**Figure 2:**
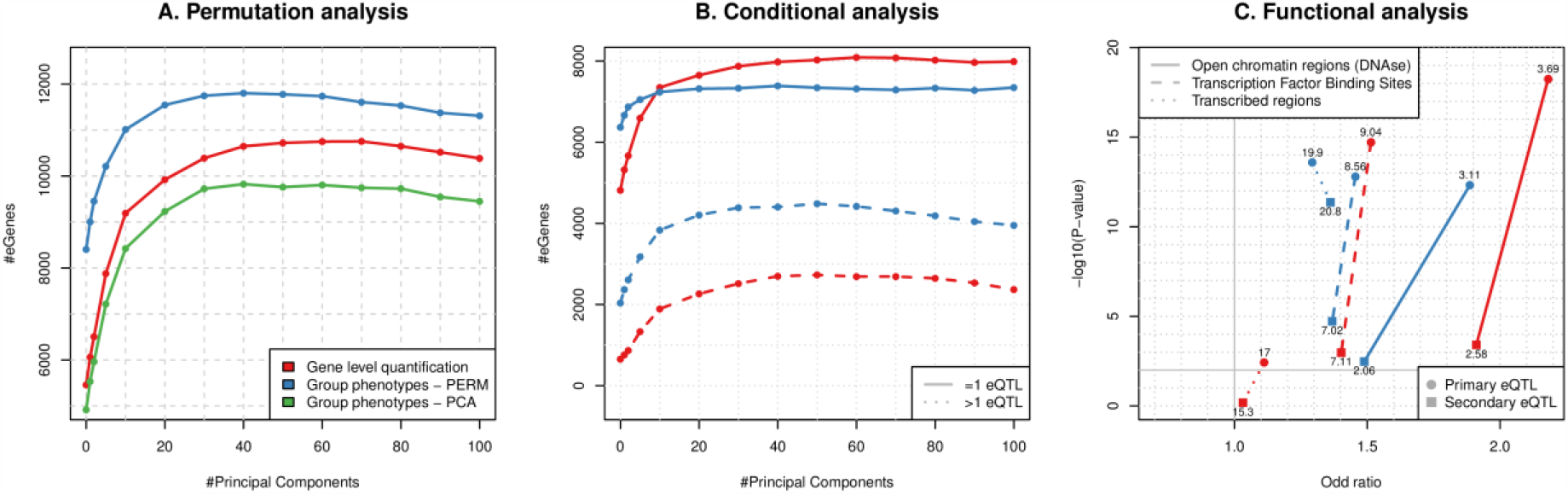
Outcome of the permutation, conditional and functional analyses on Geuvadis. Panel **(A)** shown the number of eGenes discovered (y-axis) as a function of the number of Principal Component (x-axis) used to correct for technical variance for three different ways of aggregating signal at multiple exons:at the quantification level (in red) or at the QTL mapping level (supplementary method 7) by using ethier the extended permutation scheme (in blue) or Principle Component Analysis (in green). Panel **(B)** shows the numbers of eGenes (y-axis) with a unique eQTL (solide lines) or multiple eQTLs (dotted lines) as a function of the number of Principal Components (x-axis) used to correct for technical variance. This is shown for two approaches for aggregating the signal at multiple exons: at the quantification level (in red) or at the QTL mapping level by using the extended permutation scheme (in blue). Panel **(C)** shows the enrichments of the 4 types of eQTs resulting from the analysis performed for panel (B) (primary versus secondary eQTLs and gene quantification versus phenotype grouping) within 3 types of functional annotations (supplement 12.2). The odd ratios and the -log10 of the enrichment P-values are shown on the x-axis and y-axis, respectively. The percentages of eQTLs falling within these annotations are shown next to the corresponding points.

Furthermore, QTLtools can also perform conditional analysis to discover multiple proximal molQTLs with indpendent effects on a molecular phenotype. To do so, it first uses permutations to derive a nominal p-value threshold per molecular phenotype that varies and reflects the number of independent tests per *cis*-window. Then, it uses a forward-backward stepwise regression to (i) learn the number of independent signals per phenotype, (ii) determine the best candidate variant per signal and (iii) assign all significant hits to the independent signal they belong to (supplementary method 8). We applied this conditional analysis on Geuvadis and discovered that ˜38% of the significant genes have actually more than one eQTL (figure 2B); some of them having up to 6 independent eQTLs (supplementary figure 13). Interestingly, we also find that combining the conditional analysis with the phenotype grouping approach described above could help to discover even more signals (figure 2B). We confirm that the new discoveries resulting from theses analyses in Geuvadis have high replication rates within an independent data set (GTEx [4]) suggesting that these are discoveries (supplementary method 9, supplementary figure 14).

Beyond mapping proximal molQTLs, QTLtools also includes methods to discover distal (i.e. *tran*-acting)molQTLs. The first method we implemented relies on permuting all phenotypes together in order to drraw from the null distribution of associations while preserving the correlation structure within genotype and phenotype data intact (supplementary method 10.1). By repeating this permutation scheme multiple times (e.g. 100 times in our experiments), we can obtain empirically calibrated Quantile-Quantile plot that properly shows signal enrichment (supplementary figure 15) and can estimate the False Discovery Rate (FDR) for all the most significant associations:in Geuvadis, we could find 52 genes with at least one significant signal in trans at 5% FDR. Given th ermutation scheme is computationally intensive (˜450 CPU hours for 100 permutation),we also designed an approximation of this process that gives reasonably close FDR estimates while being multiple orders of magnitude faster (˜7 CPU hours; supplementary method 10.2). Given that the whole genome is effectively tested for each phenotype, we quickly build a null distribution of associations for a single phenotype by permutations. We then use this null distribution to adjust each nominal P-value for the number of variants being tested and then standard FDR methods [12] on the resulting set of adjusted p-values to correct for the multiple phenotypes being tested. In practice, this approach can be seen as an extension in *trans* of the mapping strategy we use in *cis* and gives FDR estimates that are close to those obtained with the full permutation pass (supplementary figure 16) while being way faster to obtain (˜64 times faster in our experiments).

Finaly we also implemented multiple methods to integrate collections of molQTLs with two types of external data: functional genome annotations and GWAS results. First, QTLtools can estimate if a molQTL and a variant of interest (typically a GWAS hit) pinpoint the same underlying functional variant. To do so, it uses Regulatory Trait Concordance (RTC; supplementary method 11) [13];a sophisticated conditional analysis scheme designed to account for Linkage Disequilibrium (LD)as a confounding factor when co-localizing moiQTLs and GWAS hits. This can be used, for instance, the subset of GWAS hits that are likely mediated by molQTLs; a useful piece of information to understand the function of GWAS hits. When applied on Geuvadis and the NHGRI-EBI-GWAS catalog [14], we estimated to which extend the disease associated variants reported in this catalog overlap with eQTLs (supplementary figure 17). Alternatively, QTLtools can also look at the overlap between sets of molQTLs and functional annotations as those provided by ENCODE [8].Specifically,it can compute the density of annotation around molQLS locations (supplemeantary mathod 12.1) and,when they do overlap,estimate if it is more often than what is expected by chance (supplementary mathod 12.2).This basically allows inspecting visually and statistically the distribution of functional annotations around molQTLs. When using this on the various sets of eQTLs we discovered so far, we find that they tend to fall within transcription factor binding sites and open chromatin regions (supplementary figure 18), in line with previous knowledge on eQTLs [1].

All the functionalities described above have been implemented in C++ for high performance, in a modular way to facilitate future implementation of additional functionalities by the community.In additional,QTLtools has been designed so that the computational load can be easily distributed across the multile CPU cores typically available on a compute cluster. The set of tasks that require to be run on a per individual basis (e.g. QC the sequence data) are straightforward to parallelize:a compute job per individual. For population-based tasks, such as QTL mapping for example,the input data is split into small genomic chunks that can be run conveniently and independently on distinct CPU cores. In practice, this allows running all the experiments described above in relatively short running times (table 1), so that the full set of analyses described above can be performed in ˜1,327 CPU hours (=˜55 CPU days).

QTL tools the first software package that integrates all functionalities required to easily and rapidly go from the raw sequence and genotype data to reliable collections of proximal and distal molecular QTLs.It includes multiple new and powerful statistical methods to prepare and control the quality of the data, to map proximal and distal QTLs and to finally integrate those with GWAS results and functional annotations. It also offers a unique framework for the community to develop further additional methods or alternative to the ones already included, so that molecular QTL analysis can be more seamless among laboratories. By its integrative design and efficient implementation,QTLtools decreases drastically the time needed to set up and run the various analysis pipelines traditionally needed by molecular QTL studies, therefore allowing researchers to spend more efforts on the interpretation and validation of their results.

## Author contribution

OD, HD, AAB and ED designed the research. OD and HO implemented the methods. OD analyzed data. AF and NP helped to test the various functionalities. OD and ED supervised the research. OD wrote the paper.

## Acknowledgment

This research is supported by grants from European Commission SYSCOL FP7, European Research Council,Louis Jeantet Foundation, Swiss National Science Foundation, SystemsX, the NIH-NIMH (GTEx) and Helse Sør Øst. The computations were performed at the Vital-IT Swiss Institute of Bioinformatics.

